# Female cichlids attack and avoid—but will still mate with—androgen receptor mutant males that lack male-typical body coloration

**DOI:** 10.1101/2023.11.02.565323

**Authors:** Megan R. Howard, Maxximus G. Ramsaroop, Andrew P. Hoadley, Lillian R. Jackson, Mariana S. Lopez, Lauren A. Saenz, Beau Alward

**Author notes:** These authors share first authorship (Megan R. Howard, Maxximus G. Ramsaroop).

## Abstract

A key challenge in animal behavior is disentangling the social stimuli that drive conspecific behaviors. For behaviors like birdsong, insights can be made through the experimental isolation of relevant cues that affect behavior. However, for some species like teleost fish, putative sexual signaling cues are inextricably linked to others, making it difficult to parse the precise roles distinct signals play in driving conspecific behaviors. In the African cichlid *Astatotilapia burtoni*, males are dominant or subordinate, wherein bright coloration and territorial and courtship behavior inextricably correlate positively with rank. Here, we leveraged androgen receptor (AR) mutant male *A. burtoni* that lack dominance-typical coloration but not behavior to isolate the role of male coloration in driving female mating behaviors in this species. We found in independent behavioral assays that females behave aggressively towards AR mutant but not WT males but still mated with both types of males. Females showed enhanced activation of *esr2b+* cells in the hypothalamus when housed with either mutant or WT males and this activation scaled with spawning activities. Therefore, there is not a simple relationship between male coloration and female mating behaviors in *A. burtoni*, suggesting independent sensory mechanisms converge on hypothalamic *esr2b+* cells to coordinate behavioral output.

## Introduction

A key goal in animal behavior and sensory biology research has been to disentangle the specific sensory stimuli that control behaviors across species [1]. Concerning reproductive behaviors, much focus has historically been given to the role of male sexual signaling systems in driving female mating behavior. Decades of research have revealed that auditory cues, chemosensory signals, behavioral displays, coloration, and hormonal factors are all known to impact female mate choice in fish, reptiles, insects, birds, mammals, and humans [2–5].

Identifying the role of specific male signals in guiding female mating behaviors, however, can be challenging. For instance, if variation in signaling traits along multiple modalities correlate with female mating responses, studying the role of individual signaling traits may prove difficult. Thus, precisely identifying the roles of distinct sensory modalities in modifying behavior is challenging Therefore, a major goal of research into the influence of different signaling modalities affecting conspecific behavior has been to experimentally isolate variation only in a single trait of interest.

*Astatotilapia burtoni*, an African cichlid fish, develop plastic social hierarchies in which an individual male’s social status is reflected by his body coloration and behavior [6–8]. Dominant males are brightly colored with a dark eye bar, defend a territory, and mate frequently, while non-dominant males are drably colored, do not defend a territory, and do not mate. The unique ability of male *A. burtoni* to quickly alter the intensity of their coloration based on their social status or environment makes these fish a fascinating model for exploring the multifaceted systems controlling social behavior and mate preference [9–12]. *A. burtoni* are a mouthbrooding species whose courtship ritual involves a male performing a suite of behaviors that culminate in spawning and the fertilization of eggs [13]. First, the male will chase the female; then he will perform a quiver towards the female, in which he arches his body, displays egg spots on his fin, and rapidly vibrates his whole body. Following quivers, the male will typically lead the female back to his territory. If she follows, this may result in circling behaviors where the male and female peck at each other’s anogenital regions to stimulate the release of gametes. The female scoops up her released eggs into her mouth, which are fertilized by sperm the male releases into the water. This process in females is thought to be initiated and mediated by multiple sensory cues from the male, including chemosensory signals, auditory cues, visual cues, social status, coloration, and behavior [14,15].

One of the most striking male features that has been assumed to guide female behavior is the bright coloration displayed by dominant males. However, no study has provided evidence to support the hypothesis that male coloration mediates female mate preference in *A. burtoni*. This is because coloration correlates tightly with all aspects of social status, which can change rapidly. Indeed, within minutes after being isolated, an initially non dominant male will begin to express vibrant colors and behaviors typical of a dominant male [8,10,16,17]. No study has provided evidence to support the hypothesis that male coloration mediates female mate preference in *A. burtoni*. This is because the only way to mute body coloration in *A. burtoni* is to socially suppress the male, which suppresses other aspects of social status, including behavior. Therefore, current experimental manipulations in male *A. burtoni* do not allow for an accurate interpretation of the role of coloration in guiding female mating behaviors given the inextricable confounds of behavioral changes associated with social status suppression.

Using androgen receptor (AR) mutant *A. burtoni* engineered using CRISPR/Cas9 gene editing, we have previously shown that distinct AR genes are required for non-overlapping traits of social dominance in males such as coloration, behavior, and reproductive physiology [18]. ARβ mutant males perform normal levels of mating and territorial behaviors, but lack the bright yellow or blue body coloration typical of wild-type (WT) dominant males. This study aimed to leverage ARβ mutants to isolate male body coloration from male typical behavioral patterns and explain how male coloration impacts female mate preference in *A. burtoni*.

## Methods & Materials

### 1.1 Animals

Adult *A. burtoni* were bred from a wild-caught stock that originated from Lake Tanganyika, Africa and housed in environmental conditions that mimic their natural equatorial habitat (28 °C; pH 8.0; 12:12 h light/dark cycle with full spectrum illumination and constant aeration). Aquaria contained gravel-covered bottoms with terra cotta pots cut in half to serve as shelters. Fish were fed cichlid flakes daily. All experimental procedures were approved by the University of Houston Institutional Animal Care and Use Committee (Protocol #202000001).

### 2.1 Dichotomous Mate Choice Assays

Females were raised and maintained in mixed-sex community tanks until the day of experimentation. Gravid (egg bearing), sexually mature females were chosen from these community tanks in the morning after being fed and transferred to the experimental tank (Fig. 1A). The experimental tank design was inspired by similar designs for other female preference assays [19–21]. A 37.8-liter rectangular experimental tank was fully covered by light brown plastic placed along the inside walls of the tank to prevent fish from seeing outside the tank or seeing their own reflection. The three compartments were separated by two PDLC smartglass (SW, 29.7 cm x 21.1 cm) borders that could be remotely activated using a smart plug (Amazon Alexa) to become transparent or opaque as needed and allowed for the free flow of water between compartments (Fig. 1A). Experiments were remotely recorded from overhead using a Wifi-enabled camcorder (Canon VIXIA HF R80). This setup also included overhead dimmers (Neewer) and four soft yellow light lamps (Bohon) at each side.

**Figure 1.**
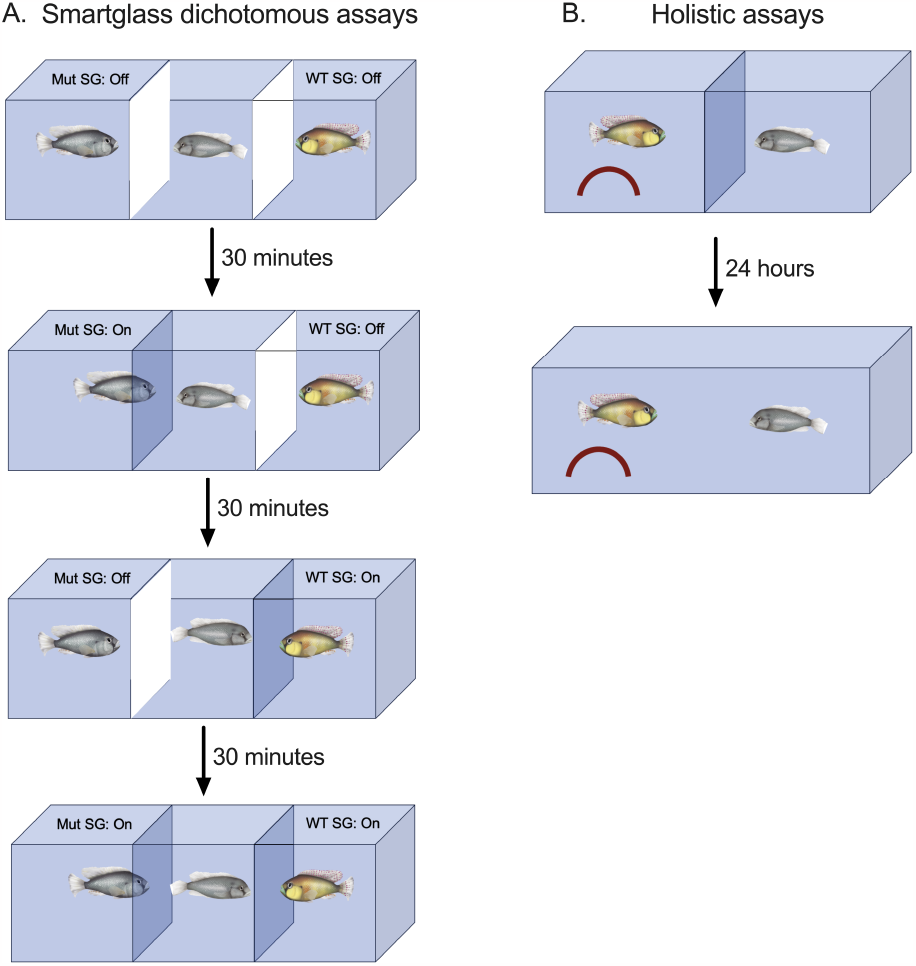
Illustration of the behavioral assays used. A) We used smartglass (SW, PDLC; 29.7 cm x 21.1 cm) to control visibility of each fish for the dichotomous choice assays. The borders were placed on both sides of the female, and a Mut or WT male were on either side. White borders in the illustration indicate opaque smartglass borders. We randomized the order in which each female saw through a Mut or WT smartglass. B) In the holistic assays, a male and female were placed into a tank in which they were separated by a transparent, perforated plexiglass barrier. 24-hours later, the barrier was removed and their behaviors recorded for 30 minutes.

At the beginning of a trial, a WT male is randomly placed in the three-compartment experimental tank’s left or right compartment. Concurrently, an ARβ mutant (“Mut”) male is placed in the empty compartment on the opposing end. The female is placed in the center compartment. Smartglass borders prevent the female and males from seeing each other during the initial 30-minute acclimation period of the assay (this last bit is confusing and should be mentioned in the next sentence). The assay is made up of four phases that proceed in the following order: 1) the video recording begins immediately after all 3 subjects are placed in the tank; the smart-glass borders remain opaque for 30 minutes to allow the fish to acclimate to a new environment; 2) after the first 30 minutes, either the right or left smartglass border (selected using a random number generator) is activated and becomes transparent, while the opposing border remains inactive; 3) at the 1 hour mark, the previously activated border is deactivated and the previously inactive border is activated; and 4) after 1 hour and 30 minutes, both borders are activated. At the end of the final 30-minute phase, the female was immediately removed and euthanized in ice-cold water before body measurements were taken and cervical transection and tissue collection was performed. The males were returned to their respective tanks until the next trial. The experimental tank was washed and air dried for approximately 24 hours between trials and 10 trials were performed in total using n=10 females and n=2 stimulus males (1 WT and 1 Mut male).

### 2.2 Holistic Mate Choice Assays

Based on the results of the dichotomous choice assay (see “Results” section), we decided to measure how females behave with either a WT or Mut male in a manner that allowed holistic behavioral interactions with one male at a time (Fig. 1B). Females were raised in community tanks, then housed over the duration of the experiment in a tank with several females and a WT male that was visibly and chemically accessible but physically isolated to prevent spawning. We ran 24-hour assays in which one of the males was added to a non-recirculating assay tank across a permeable, transparent border from a female. Each female was selected from the community tank based on the visual appearance of gravidity. A male was placed on the left side of the permeable plexiglass border along with a half terracotta pot that could be used as a spawning site. Then, the female was placed on the right side of the border by herself. We allowed the fish to sit in the assay set up for approximately 24 hours to acclimate to each other. After 24 hours, we recorded their behavior for 30 minutes, then removed the border between them, and recorded the first 30 minutes of their interaction. Immediately after these 30 minutes of interaction, the female was removed and euthanized in ice-cold water before body measurements were taken and cervical transection and tissue collection were performed. Two WT and two Mut males were used as stimulus males. These males were housed in a separate room from the females, each in their own tank along with two stimulus wild type females each. The order of the inclusion of a male in an assay was randomly determined. Final sample sizes for each group were: Females with WT: n=6; Females with Mut: n=6; we also included females that were placed into the tank without a male on the other side of the border, a condition referred to as “Empty”: n=6.

### 2.3 Body measurements and tissue collection

Immediately following euthanizing in ice-cold water, standard length (SL) and body mass (BM) were recorded, blood was collected from the caudal vein, and then rapid cervical transection was performed. The brain was collected and immediately transferred to 4% paraformaldehyde (PFA) of pH=7.0. The gonads were then weighed. For fish that released their eggs during the assay and began mouthbrooding, we removed those eggs and weighed them as a proxy for gonad weight. Gonad mass was expressed as % body mass. 24 hours after the brain was placed into PFA, it was moved into 15% sucrose; after another 24 hours it was moved to 30% sucrose. After 24 hours in 30% sucrose, the brain was embedded in Neg50 and stored at –80 degrees C. The brains of females from the holistic assay were collected for measuring gene expression and were sectioned at 30 μm into three series and mounted onto Superfrost plus slides, dried for 24 hours, and stored at –80 degrees C until they were used for HCR.

### 2.4 Behavioral Scoring

#### 2.4.1 Noldus Ethovision XT Behavioral Scoring

The analysis of the dichotomous mate choice assays was performed using Noldus Ethovision XT Version 14. Using Ethovision, the dichotomous mate choice assays were separated into three arenas tracking the central female and both males across their respective arenas. Subsections or “Interest Zones”, were drawn along the length of the left and right side of the central arena indicating time spent near either the WT or ARβ mutant in the adjacent arena. These “Interest Zones” were made to be approximately 1 body length in width, matched to the body length of the female in each trial. The frequency of visits (count) and cumulative duration(s) of each visit were measured for each zone in each trial. Time spent in interest zones and frequency of visits were used in tandem with individual behavioral scoring to evaluate if females showed an interest in either WT or ARβ mutant males. Heatmaps showing time spent in areas of the experimental tank for each fish were generated for each trial.

#### 2.4.2 BORIS Behavioral Scoring

Individual behaviors were scored using BORIS (Behavioral Observation Research Interactive Software), a manual behavioral scoring software. Both dichotomous mate choice assays and holistic behavioral assays were recorded for the full duration of the trial then scored. Table 1 lists the behaviors scored, their respective definitions, and the broad categories used to interpret the function of said behaviors.

**Table 1.** Ethogram used for scoring behaviors in BORIS.

| Behavior | Description | Category |
| --- | --- | --- |
| Approach | Fish A (M/F) approaches Fish B(M/F) slowly in direct path and stops within 0.5 body lengths of B | Exploratory |
| Attack | Fish A accelerates towards Fish B, making contact and vigorously wriggling body | Aggression |
| Bite | Fish A approaches Fish B and attempts to lightly bite at the anal fin of Fish B | Reproductive |
| Bore | Fish vigorously wriggles body while contacting its mouth with the glass for more than 3 seconds and traveling a distance of at least 1 body length in any direction. | Exploratory/General Motivation |
| Chase Female | Male quickly moves towards female (coupled with reproductive behaviors) | Reproductive |
| Flee | Fish quickly turns away from its initial orientation and moves away in one swift direct action. | Avoidance |
| Graze | Fish swims back and forth along glass/border horizontally or vertically while continuously contacting the surface with its mouth for more than 3 seconds | Stress |
| Lateral Display | while lateral to Fish B, Fish A conforms body in a concave fashion several times in close vicinity to fish B | Aggression |
| Lead Swim | male turns and swims away from female accompanied by back-and-forth motions of the tail (waggles), signaling to the female to follow it | Reproductive |
| Pot Entry | Fish enters terracotta pot | Reproductive |
| Quiver | Fish A vibrates body when near Fish B, body conforms to a convex shape; fish will typically remain in the same spot during event | Aggression |
| Retreat | Fish A suddenly swims backwards a short distance while facing original direction away from Fish B | Avoidance |
| Rostral Display | Fish A flares opercula at Fish B. | Aggression |
| Spawn | Both fish perform spawning ritual consisting of bites and circling each other | Reproductive |
| Short Flee | Fish quickly turns away from its initial orientation and moves away in one swift direct action, moving less than 1 full body length away | Avoidance |
| Tap | Fish taps glass/border gently by making contact with mouth | Exploratory |

### 3.1 Hybridization Chain Reaction

HCR was performed according to manufacturer protocols, recommendations, and reagents, unless otherwise noted. In summary, on day 1 of the protocol, tissue slides were exposed to LED light for 60mins to reduce autofluorescence. Subsequently, tissue was immersed in 0.2% Triton X-100 (in 1x PBS) (Thermo Fisher; Gibco) for 30 minutes to increase signal detection. Next, tissue was dehydrated with serial concentrations of EtOH (Thermo Fisher) at 50%, 70%, and 100%, before adding Proteinase K (1:2000) (Thermo Fisher) in 1xPBS. HCR probes were then hybridized to tissue by adding 16nMs of RNA probe in hybridization buffer for 12-16hrs at 37°C. The HCR probes used against *esr2b* and *egr1* were engineered by Molecular Instruments. Following the hybridization step, specific probe amplifiers tagged with distinct fluorophores were added at 60nM into a single amplifier solution. Slides were incubated in amplifier solution at room temperature in a dark chamber overnight. B3-488 amplifier was used to detect the *egr1* probe, while B2-647 amplifier was used to detect the *esr2b* probe. Slides were then washed in 5X SSCT (0.1%Tween-20) (Gibco; Thermo Fisher), and cover slipped using Prolong Gold (Invitrogen) mounting media with DAPI. Tissue from one fish from each group suffered from very poor tissue quality and adhesion, so these fish were not included in cell quantification. Final sample sizes for cell quantification: WT: n=5; Mut: n=5; and Empty: n=5.

### 3.2 Quantification of egr1 & esrb2

Slides were imaged using a Nikon Eclipse 80i Microscope MicroFireTM at 20x magnification in YFP, cy5, and DAPI wavelength filters in the ventral part of the ventral telencephalon (Vv) (lateral septum homolog) and the preoptic area (POA) using multichannel, multiplane image capture. Our goal was to quantify the amount of *esr2b*+ cells that were active as measured by co-expression with *egr1*. To do this, we used Fiji ImageJ cell counting software, counting the cells in which *esr2b* was present and then which cells expressed both *esr2b* and *egr1*. We counted two subsequent sections per brain region with the highest *esr2b* expression for analysis. *Esr2b+* expression as indicated by a clear cy5 signal was found to be in the nucleus or the cytoplasm. *Esr2b+* cells that co-expressed *egr1* were determined by a cy5 signal in the nucleus or the cytoplasm that also exhibited a ring-like pattern of YFP expression. We determined using z-stacks that co-expression occurred consistently based on these signal characteristics.

### 4.1 Statistical Analysis

A two-way repeated measures ANOVA was used with male genotype (WT or Mut) and smartglass condition as the within-subjects factors influencing female behavior with a Greenhouse–Geisser correction. Female or male behaviors during the holistic dyadic interactions were analyzed statistically using unpaired t-tests or Mann Whitney tests when normality and equality of variance assumptions were not met. A Fisher’s exact test was used to test the difference in the frequency of females performing a lateral display at WT or Mut males. One-way ANOVAs were used to test the effects of male social condition (Empty, WT, Mut) on percentage of *esr2b+* cells that expressed *egr1+* in the Vv and POA. Following a significant Omnibus ANOVA factor, post-hoc Tukey’s tests were used for multiple comparisons. Pearson’s correlation analyses were conducted between mating related behaviors and percentage of *esr2b+* cells that expressed *egr1+* in the Vv and POA. Effects were considered significant at *P*≤0.05.

## Results

### Females visits transparent border regardless of genotype

Heatmaps showing movement patterns during each smartglass condition suggested females spent most of their time near whichever male had a transparent smartglass (Fig. 2 A-D). Males showed a similar pattern wherein they spent their time near the smartglass when it was transparent. When both of the smartglasses were on, rendering both males visible, heatmaps showed that females varied considerable in which male they spent their time near and in some cases, they spent a relatively equal amount of time near both males.

**Figure 2.**
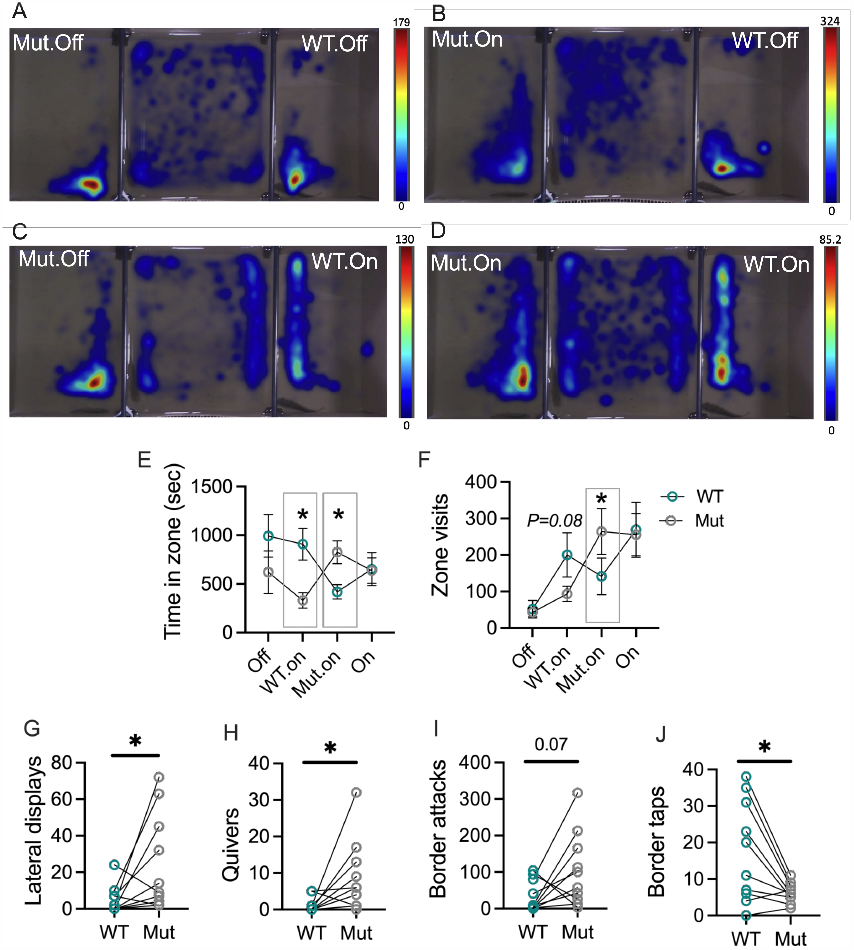
Females spend time near WT and Mut males but behave differently towards them. A-D) Heat maps overlaid on representative conditions within a given trial from a female with two flanking males. Scale bars were calibrated according to the longest time spent in one area for that trial. Coloration of an area indicates fish spent time in that area progressing from blue to red in hue as more time is spent in the area. E) Females spent more time in the WT zone when its smartglass was on and the Mut smartglass was off and vice versa. F) Females visited the Mut zone significantly more times when the Mut smartglass was on and the WT smartglass was off and the opposite comparison did not reach statistical significance. G-J) Females performed different behaviors towards WT and Mut males depending on whose smartglass was on. *=*P<0*.*05*. E-F) Circles represent the Mean±Standard Error of the Mean (SEM). G-J) Circles connected by straight lines represent individual females.

Recording during the both-off condition for one trial malfunctioned and thus there are no behavioral measures for this female in this condition. A mixed-effects model was thus applied that could handle a missing value. A two-way repeated measures ANOVA with the both-off condition completely removed yielded similar results to that of the mixed-model. Results of the mixed-model indicated a significant effect of condition*genotype on Time in zone (smartglass condition: F_(1.93, 17.34)_ = 0.72, *P* = 0.4964; genotype: F_(1.00, 9.00)_ = 1.58, *P* = 0.2410); smartglass condition*genotype: F_(1.96, 16.36)_ = 4.43, *P* = 0.0295. Specifically, females spent more time in the WT zone when its smartglass was on and the Mut smartglass was off (Fig. 3E; Tukey’s comparison, *P* = 0.0224) and vice versa (Fig. 2E; Tukey’s comparison, *P* = 0.0446). A similar pattern was observed for visits to zone (smartglass condition: F_(1.37, 12.31)_ = 9.99, *P* = 0.0050; genotype: F_(1.00, 9.00)_ = 0.02, *P* = 0.8955); smartglass condition*genotype: F_(1.91, 15.95)_ = 6.79, *P* = 0.0079). Specifically, when the Mut smartglass was on, females visited the Mut zone significantly more than the WT zone (Fig. 2F; Tukey’s comparison, *P* = 0.0154). When the WT smartglass was on, females visited the WT zone more than the Mut zone, but this difference did not reach statistical significance (Fig. 2F; Tukey’s comparison, *P* = 0.0883).

**Figure 3.**
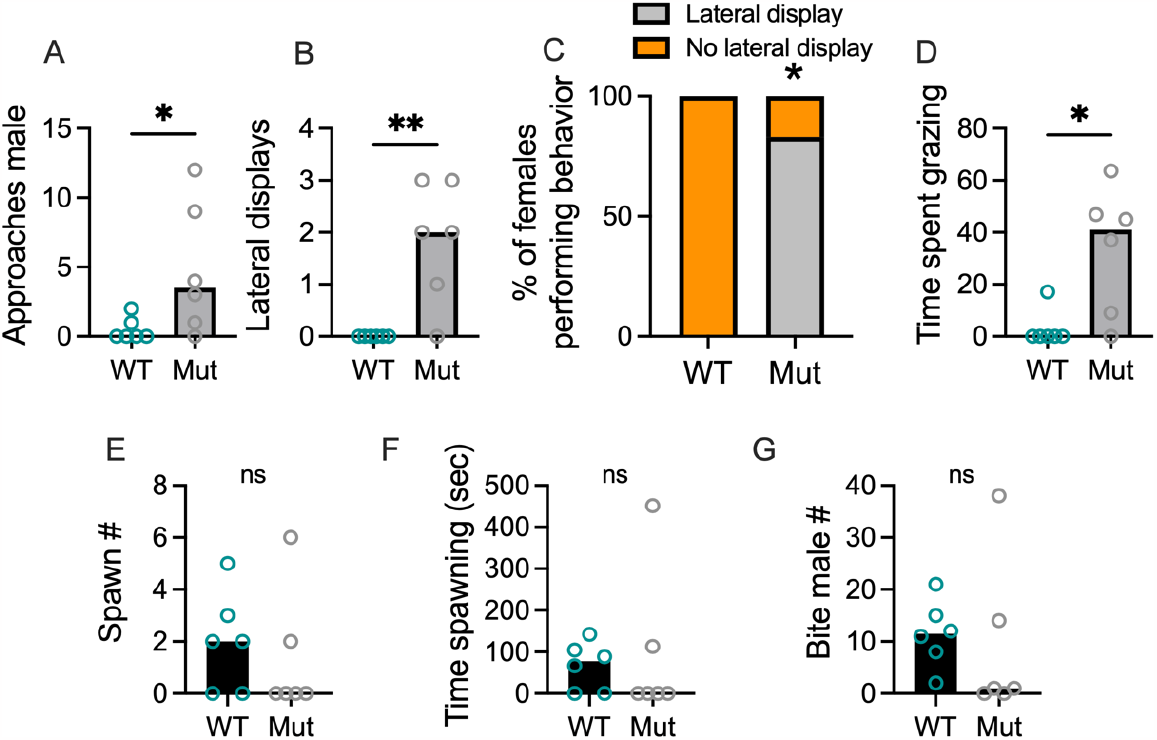
In holistic trials, females performed different behaviors towards WT and Mut males, they still spawned with both types of males. A-C) Females approached and performed aggressive displays towards Mut males more than WT males. D) Grazing, which may reflect an anxiety of avoidance state, was done for more time by females housed with Mut males compared to WT males. E-F) There were no significant differences in spawning behaviors. G) Biting, which can occur before and during spawning, did not differ. *=*P<0*.*05*. **=*P<*0.01. Circles represent individual females. Bars show the Mean.

### Females behave more aggressively toward Mut than WT males when both are simultaneously visible

Since females did not differ in the number of visits to both WT and Mut zones, we wondered whether they performed different behaviors towards either male when in their zones. We found that females performed more lateral displays (Fig. 2G; t_9_=2.3, *P* = 0.0479) and quivers (Fig. 2H; t_9_=2.4, *P* = 0.0421) towards Mut males compared to WT males. Similarly, performed more border attacks at Mut males compared to WT males, but this effect was not significant (Fig. 2I; t_9_=2.0, *P* = 0.0731). Females performed significantly more taps at WT males compared to Mut males (Fig. 2J; t_9_ =3.1, *P* = 0.0134). These findings indicate that while females visit WT and Mut zones at similar rates, they perform different behaviors.

Given our findings on female behaviors towards either male, we queried whether behaviors performed by the males related to that performed by the females. We compared WT and Mut male positional data across the both-off and both-on smartglass conditions as well as the condition during which only their respective smartglass was on. There was a significant impact of genotype but not condition or a genotype by condition interaction (smartglass condition: F_(1.04, 18.65)_ = 0.1993, *P* = 0.6692; genotype: F_(1.00, 18.00)_ = 6.64, *P* = 0.0190; smartglass condition*genotype: F_(2, 36)_ = 0.75, *P* = 0.4807) where Mut males spent significantly more time in the female zone compared to WT males (Fig. S1A). We observed a significant effect of smartglass condition but not genotype or a smartglass condition by genotyope interaction on vists to the female zone (smartglass condition: F_(1.53, 27.48)_ = 5.987, *P* = 0.0114; genotype: F_(1.00, 18.00)_ = 2.75, *P* = 0.0047; smartglass condition*genotype: F_(2, 36)_ = 0.008, *P* = 0.9925), where visits to the female zone increased when both smartglasses were on or the respective smartglass was on (Fig. S1B; Tukey’s tests: Both-off versus Both-on, *P* = 0.0327; Both-off versus On, *P* = 0.0306; Both-on versus On, *P* = 0.4228). Thus, in addition to females performing more aggressive behaviors towards Mut males than WT males, Mut males spend significantly more time in the female zone than WT males.

We also scored state behaviors of the WT and Mut males for all trials during the condition in which both smartglasses were on. Mut males performed significantly more bores against the female smartglass (Fig. S2A; *U=*21.5, *P* = 0.0297), a measure of exploratory behavior that may be related to motivation to make it across the barrier. WT and Mut males did not differ in the number of grazes performed (Fig. S2B; *U=*39.5, *P* = 0.4331) nor did they differ on any mating related behaviors (Quivers: Fig. S2C; *U=*34.0, *P* = 0.2310; Lead swims: Fig. S2D; *U=*35.5, *P* = 0.2809). Mut males tended to perform more lateral displays than WT males, an effect that did not reach statistical significance (Fig. S2E; *U=*26.0, *P* = 0.0573). WT and Mut males did not differ in the number of attacks at the female smartglass (Fig. S2F; t_18_=1.7, *P* = 0.0964). Therefore, while WT and Mut males did not differ in the mating behaviors they performed during the holistic assay, Mut males may have been more motivated to get across the female border.

### In a holistic assay, females perform lateral displays at and try to avoid Mut males, but will still mate with them

Our observations that females visited Mut males but perform aggressive behaviors towards them led us to wonder whether females would mate with Mut males during an assay in which they were allowed both visual and physical access. To test this, we performed holistic behavioral assays in which a single female could interact with an individual Mut or WT male (Fig. 3). Females performed more approaches (Fig. 3A; *U=*5.5, *P* = 0.0455) and lateral displays (Fig. 3B; *U=*3, *P* = 0.0152) towards Mut males than WT males. A Fisher’s exact test revealed the frequency of females performing a lateral display towards a Mut male was higher than that towards a WT male (Fig. 3C; 0/6 performed a lateral display towards WT males; 5/6 performed a lateral display towards a Mut male; *P* = 0.0152). Females also spent more time grazing along the tank edges, a potential reflection of stress or attempts to escape, when housed with Mut males compared to WT males (Fig. 3D; *U=*4.5, *P* = 0.0281). Surprisingly, females spawned with both WT and Mut males (Video S1 and Video S2; 4/6 females spawned with WT males; 2/6 spawned with Mut males; Fisher’s exact test: *P*=0.5671) and did not significantly differ on any measure of mating behaviors (Fig. 3E-G; Spawn #, t_10_=0.5, *P* = 0.6072; Time spent spawning, *U=*15, *P* = 0.6212; Bite male #, t_10_=0.4, *P* = 0.7183). Thus, while females perform aggressive displays and exhibit avoidance or stress responses while housed with Mut males, they still mate with them.

We also scored behaviors performed by the males during the holistic trials. Mut males performed significantly more bores and grazes compared to WT males (Bores: Fig. S3A; *U=*0, *P* = 0.0022; Grazes: Fig. S3B; *U=*5.5, *P* = 0.0411). On the other hand, WT males performed significantly more quivers, chases, and lead swims than Mut males (Quivers: Fig. S3C; t_10_=2.8, *P* = 0.0173; Chases: Fig. S3D; t_10_=3.0, *P* = 0.0130; Lead swims: Fig. S3E; *U=*4.5, *P* = 0.0303). WT and Mut males did not differ in the number of lateral displays or attacks directed towards females (Lateral displays: Fig. S3F; t_10_=1.6, *P* = 0.1403; Attacks: Fig. S3G; *U=*15, *P* = 0.6991). Therefore, Mut males may attempt few courtship behaviors, likely because they are attempting to avoid females (bores, grazes) due to the agonistic behaviors being performed by the females.

### The number of egr1+ esr2b+ cells in the Vv and POA correlates with mating behaviors

We hypothesized that females housed with Mut males would show different cellular activation patterns compared to WT males in specific cells thought to be involved in mating behaviors in teleosts. Specifically, based on recent work in medaka, in the females included in the holistic assay, we quantified the percentage of *esr2b+* cells that were active as measured by co-expression with *egr1* [22], in the Vv and POA (Fig. 4 A-F), areas with high expression of *esr2b* [23,24] that are also involved in mating behaviors [25–27].

**Figure 4.**
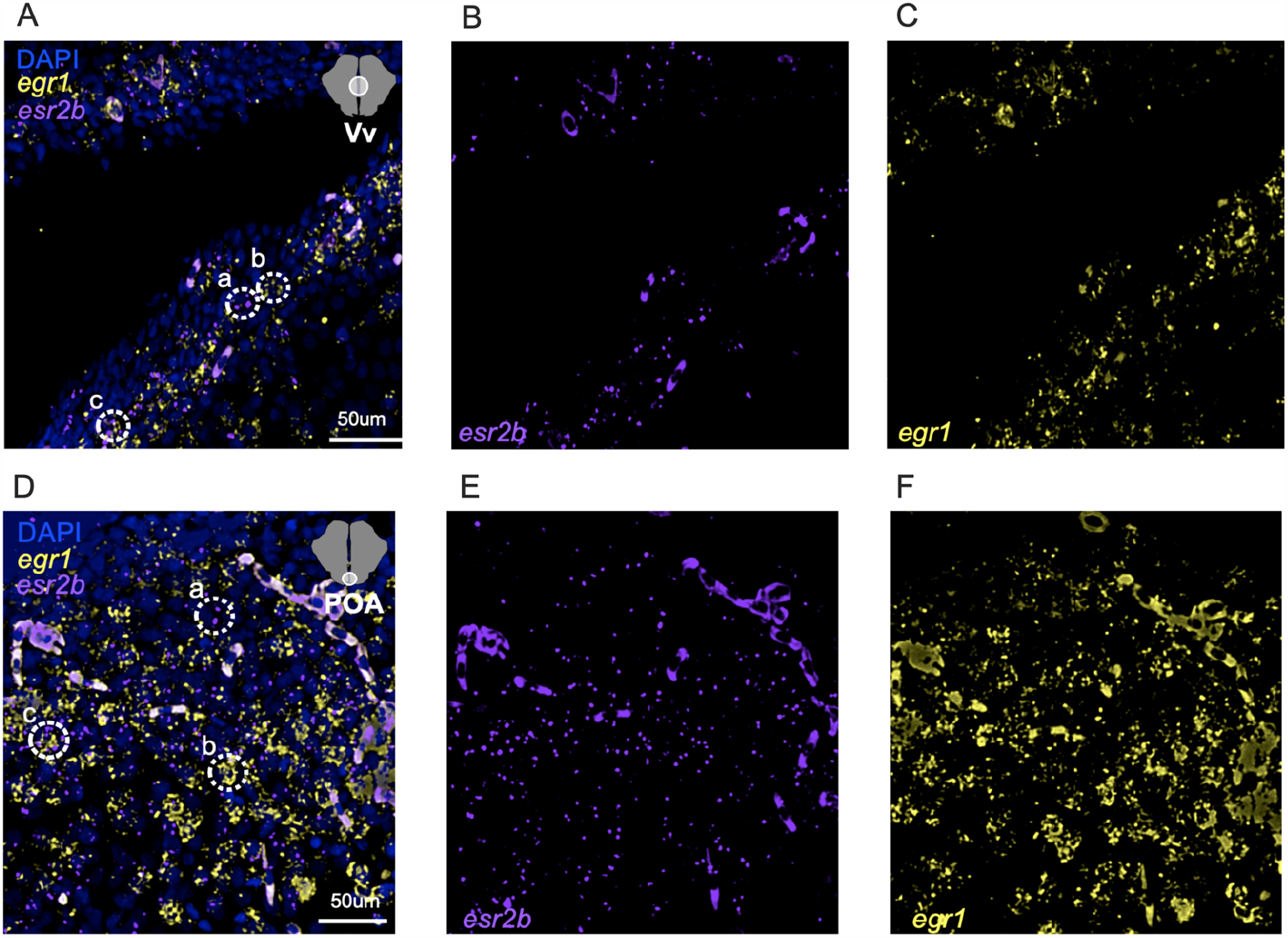
Representative photomicrographs of HCR to detect expression of *egr1, esr2b*, and co-expression. A-C) Shows the DAPI, yfp, and cy5 filters simultaneously which coincide to labels of the DAPI marker, which is a pan-cellular marker that localizes to the nucleus; *egr1*; and *esr2b*. For the Vv, in A), the dotted circle labeled “a” surrounds solely *esr2b* signal; the “b” dotted circle surrounds only *egr1;* and the “c” dotted circle surrounds a cell that expresses both *esr2b* and *egr1*. The same is true for the POA in panel D.

The effect of male housing condition (Empty, WT, Mut) on the % of egr1+ esr2b cells in the female Vv did not reach statistical significance (Fig. 5A; One-Way ANOVA, F_(2, 12)_ = 3.36, *P* = 0.0695). There was a significant effect of male housing condition on % of egr1+ esr2b cells in the female POA (Fig. 5B; One-Way ANOVA, F_(2, 12)_ = 4.34, *P* = 0.0382), wherein females housed with Mut males had a significantly higher % of egr1+ esr2b cells compared to females in the Empty condition (Tukey’s comparison, *P* = 0.0325). For both brain regions, while variation in cell % across groups was high, % of egr1+ esr2b cells in the Vv and POA of females with Mut and WT females was clearly not different, which was confirmed statistically (Vv, t_10_=0.13, *P* = 0.9021; POA, t_10_=0.87, *P* = 0.4084). We collapsed across male genotype so that females housed with males generally were compared to females housed with no male. We found females housed with a male had a significantly higher % of egr1+ esr2b cells in both regions (Vv, t_13_=2.69, *P* = 0.0185; POA, *U*=4, *P* = 0.0077). The results suggest esr2b+ neurons in the Vv and POA do not respond to aspects of the male color phenotype but instead may report the mere presence of a male in their social environment and/or reflect variation in behavioral output.

**Figure 5.**
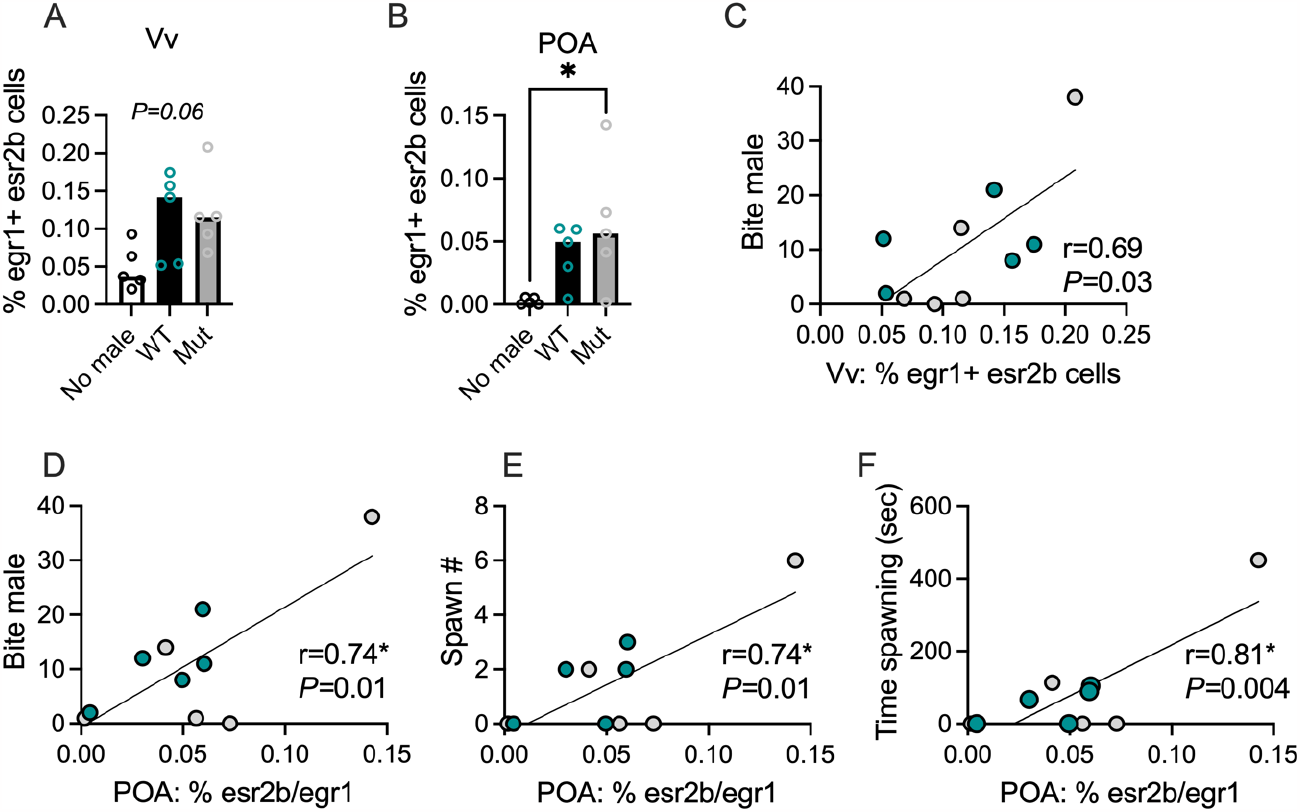
Activation of *esr2b*+ cells was higher in (A-B) females with a male and (C-F) females that performing mating behaviors. *=*P<0*.*05*. **=*P<*0.01. Circles represent individual females. Bars show Mean±Standard Error of the Mean (SEM).

To determine if the activation of esr2b+ cells relates to behavioral variation, we performed correlational analyses between the % of egr1+ esr2b cells in the Vv and POA of females housed with males to three measures of mating behaviors: bite male #, spawn #, and time spent spawning. The % of egr1+ esr2b cells in the Vv significantly positively correlated to bite male # (Fig. 5C; *r=*0.69, *P* = 0.03). In the POA, % of egr1+ esr2b cells significantly positively correlated to bite male # (Fig. 5D; *r=*0.74, *P* = 0.01), spawn # (Fig. 5E; *r=*0.74, *P* = 0.01), and time spent spawning (Fig. 5F; *r=*0.81, *P* = 0.004).

## 4. Discussion

We sought to determine whether WT male cichlids with bright, DOM-typical coloration would be preferred by females over novel AR mutant male cichlids that lack male-typical coloration. We found that the answer to this question is inordinately nuanced, prompting us to explore specific female behaviors in distinct experimental setups. Ultimately, we discovered that female *A. burtoni* behave aggressively towards males that lack coloration but will still mate with them and that the cellular population putatively involved in regulating female mating behavior was activated in females exposed to either WT or Mut males. These findings reveal that while male coloration plays an important role in guiding behavior in female *A. burtoni*, it alone is not necessary to drive female mating behaviors.

Coloration in *A. burtoni* males has been linked to physiological and behavioral differences in males [6,12,28,29] but has not been shown clearly to be used by females to guide mate preference. For instance, the most obvious connection of color to behavior is that dominant males, which are territorial and mate, are brightly colored yellow or blue while subordinate males, which are not territorial and do not mate, are drably colored [6,12]. Additionally, Korzan and colleagues found that yellow males became dominant over blue males more often than the blue over yellow [29] and dominant males are more aggressive towards other males with a color unlike their own [28]. Additionally, gravid females prefer to affiliate with dominant over subordinate males [30]. Despite this knowledge on links between color and male behavior and the preference of females to associate with dominant males, the role of male coloration in guiding female behaviors has not been definitively shown. By dissociating male color from DOM social status using ARβ mutant males that lack DOM coloration but still perform DOM behaviors, we aimed to determine whether females prefer colorful males. We predicted that there would be a clear preference by females towards colorful males as displayed by a greater amount of time in or more visits to the WT male zone over the Mut male zone in our dichotomous assay. Surprisingly, we did not observe a difference in these zone measures, as females spent an equal amount of time visiting WT and Mut males when their smartglass borders were transparent and, on average, showed no clear preference for spending time near the WT or Mut male when both smartglass borders were simultaneously transparent. Instead, we observed clear differences in behaviors performed by females towards either male, with females performing more aggressive displays towards Mut males than WT males. We predicted this difference in aggression would replicate in a holistic assay and females would not mate with Mut males. Instead, while indeed females exhibited aggression towards Mut males, they still mated with them.

We propose the following hypotheses to explain our findings. Females perform aggressive behaviors like lateral displays towards females in simulated (e.g., through the use of a mirror), dyadic same-sex social interactions, and community tanks with all females [31,32]. Given that females behave aggressively towards ARβ mutant males, we posit that females perceive Mut males as females to a certain degree. However, the results of our holistic assay and cellular quantification suggest coloration is not a sufficient trait to report unequivocally the presence of a given sex to a female. Indeed, although in the holistic assay females performed aggressive displays towards Mut males, they mated and spawned with them at a rate statistically no different to that of WT males. Therefore, the novelty of our findings lies in the fact that females behaved to ARβ mutant fish as if they were females, and *still* mated with them. This suggests the presence of independent signaling mechanisms present in females that detect multiple sensory cues, integrate them, and coordinate optimal behavioral responses. This hypothesis is in line with ideas proposed by others claiming that multiple sensory stimuli from *A. burtoni* males including behavior, coloration, pheromones, and acoustic signals affect female behavior and mating patterns [13,15,33,34]. Multiple assumptions of this hypothesis can be tested in future studies, including through the use of ARα mutant male *A. burtoni*, which produce fewer DOM-typical behaviors than WT male but still possess DOM-typical bright coloration.

Our findings on *egr1+ esr2b* cells in the Vv and POA suggest these cells may encode the motivation to mate, results that are in line with recent work from Nishiike et al [24] showing that *esr2b* is required for female-typical mating. Indeed, the percentage of *egr1+ esr2b* cells in the Vv and POA scaled positively with the number of times and time spent spawning with males regardless of the male’s genotype. Moreover, given that the percentage of *egr1+ esr2b* cells in the Vv and POA of females with either WT or Mut males were indistinguishable from each other and yet higher than that of females not housed with a male, *esr2b+* cells do not appear to be responsive to the color of males. The percentage of *egr1+ esr2b* cells in the Vv and POA did not correlate significantly with any other behaviors performed during the holistic assay, supporting the idea that these cells encode specifically the motivation to mate or an internal state related to mating. Nonetheless, future work is warranted in which the role of these cells are investigated in other contexts to precisely disentangle their functions.

Combining the current results with previous findings, we propose an overarching working hypothesis that lays the groundwork for future experiments to identify whether and how male cues guide female mating in *A. burtoni*. Our hypothesis stipulates that there exist independent mechanisms for the detection of male-typical cues that includes molecular and neural substrates that detect coloration, behavior, and pheromones that are integrated in *esr2b+* cells in the SBN and coordinate distinct behavioral responses appropriate to the specific social milieu. This hypothesis acknowledges the fact that we observed female-typical aggression directed towards males that females ultimately mated with. Each component of this working hypothesis is testable using the diverse, interdisciplinary methodological toolkit available for *A. burtoni* that includes a sequenced and annotated genome, CRISPR/Cas9 gene editing, and next-generation sequencing technologies. Studies dissecting the sensory mechanisms of mating behaviors and their underlying molecular and neural substrates in *A. burtoni* may yield novel insights into the control of social behavior broadly in teleosts and other species.

## Ethics

Experimental procedures were conducted according to the ethical guidelines for the care and use of laboratory animals. Experiments were approved by the University of Houston Institutional Animal Care and Use Committee (Protocol #202000001).

## Data accessibility

The datasets generated and/or analyzed during the current study are available in the manuscript itself, the Supplementary Information file, and online at https://github.com/AlwardLab.

## Author contributions

B.A.A., L.R.J., and M.S.L. conceived of the idea for the study; A.P.H., M.R.H. and M.G.R. designed the study; M.R.H. and M.G.R. performed the experiments; M.R.H., M.G.R., and L.A.S. collected the data. B.A.A. analyzed the data. All authors contributed to preparing and editing an earlier version of the manuscript.

## Acknowledgements

The authors thank Kathleen Munley for procedural assistance as well as Melanie Dussenne for helpful discussions on interpreting findings.

## References

1. Searcy WA, Nowicki S. 2010 Searcy, William A., and Stephen Nowicki. The Evolution of Animal Communication: Reliability and Deception in Signaling Systems: Reliability and Deception in Signaling Systems. Princeton University Press.

2. Maney DL, Cho E, Goode CT. 2006 Estrogen-dependent selectivity of genomic responses to birdsong: Estrogen-dependent genomic responses. Eur. J. Neurosci. 23, 1523–1529. (doi:10.1111/j.1460-9568.2006.04673.x)

3. McComb KE. 1991 Female choice for high roaring rates in red deer, Cervus elaphus. Anim. Behav. 41, 79–88. (doi:10.1016/S0003-3472(05)80504-4)

4. Waitt C, Little AC, Wolfensohn S, Honess P, Brown AP, Buchanan-Smith HM, Perrett DI. 2003 Evidence from rhesus macaques suggests that male coloration plays a role in female primate mate choice. Proc. R. Soc. Lond. B Biol. Sci. 270. (doi:10.1098/rsbl.2003.0065)

5. Byers J, Hebets E, Podos J. 2010 Female mate choice based upon male motor performance. Anim. Behav. 79, 771–778. (doi:10.1016/j.anbehav.2010.01.009)

6. Fernald RD. 2012 Social control of the brain. Annu. Rev. Neurosci. 35, 133–51. (doi:10.1146/annurev-neuro-062111-150520)

7. Fernald RD, Maruska KP. 2012 Social information changes the brain. Proc. Natl. Acad. Sci. 109, 17194–17199. (doi:10.1073/pnas.1202552109)

8. Maruska KP. 2015 Social Transitions Cause Rapid Behavioral and Neuroendocrine Changes. Integr. Comp. Biol. 55, 1–13. (doi:10.1093/icb/icv057)

9. Stevenson TJ, Alward BA, Ebling FJP, Fernald RD, Kelly A, Ophir AG. 2018 The value of comparative animal research: Krogh’s Principle facilitates scientific discoveries. Policy Insights Behav. Brain Sci. 5, 118–125.

10. Alward BA, Hoadley AP, Jackson LR, Lopez MS. 2023 Genetic dissection of steroid-hormone modulated social behavior: Novel paralogous genes are a boon for discovery. Horm. Behav. 147, 105295. (doi:10.1016/j.yhbeh.2022.105295)

11. Jackson LR, Lopez MS, Alward BA. 2023 Breaking through the bottleneck: Krogh’s principle in behavioral neuroendocrinology and the potential of gene editing. Integr. Comp. Biol.

12. Maruska KP, Fernald RD. 2018 Astatotilapia burtoni: A Model System for Analyzing the Neurobiology of Behavior. ACS Chem. Neurosci. 9, 1951–1962. (doi:10.1021/acschemneuro.7b00496)

13. Juntti SA, Fernald RD. 2016 Timing reproduction in teleost fish: Cues and mechanisms. Curr. Opin. Neurobiol. 38, 57–62. (doi:10.1016/j.conb.2016.02.006)

14. Maruska KP, Butler JM. 2021 Reproductive- and Social-State Plasticity of Multiple Sensory Systems in a Cichlid Fish. Integr. Comp. Biol. 61, 249–268. (doi:10.1093/icb/icab062)

15. King T, Ray EJ, Tramontana B, Maruska K. 2022 Behavior and neural activation patterns of non-redundant visual and acoustic signaling during courtship in an African cichlid fish. J. Exp. Biol. 225, jeb244548. (doi:10.1242/jeb.244548)

16. Maruska KP, Fernald RD. 2010 Behavioral and physiological plasticity: Rapid changes during social ascent in an African cichlid fish. Horm. Behav. 58, 230–240. (doi:10.1016/j.yhbeh.2010.03.011)

17. Alward BA, Hilliard AT, York RA, Fernald RD. 2019 Hormonal regulation of social ascent and temporal patterns of behavior in an African cichlid. Horm. Behav. 107, 83–95. (doi:10.1016/j.yhbeh.2018.12.010)

18. Alward BA, Laud VA, Skalnik CJ, York RA, Juntti SA, Fernald RD. 2020 Modular genetic control of social status in a cichlid fish. Proc. Natl. Acad. Sci. 117, 28167–28174.

19. Kidd MR, O’Connell LA, Kidd CE, Chen CW, Fontenot MR, Williams SJ, Hofmann HA. 2013 Female preference for males depends on reproductive physiology in the African cichlid fish Astatotilapia burtoni. Gen. Comp. Endocrinol. 180, 56–63. (doi:10.1016/j.ygcen.2012.10.014)

20. Plenderleith M, Oosterhout CV, Robinson RL, Turner GF. 2005 Female preference for conspecific males based on olfactory cues in a Lake Malawi cichlid fish. Biol. Lett. 1, 411–414. (doi:10.1098/rsbl.2005.0355)

21. Theis A, Salzburger W, Egger B. 2012 The Function of Anal Fin Egg-Spots in the Cichlid Fish Astatotilapia burtoni. PLoS ONE 7, e29878. (doi:10.1371/journal.pone.0029878)

22. Clayton DF. 2000 The Genomic Action Potential. Neurobiol. Learn. Mem. 74, 185–216. (doi:10.1006/nlme.2000.3967)

23. Maruska KP, Butler JM, Anselmo C, Tandukar G. 2020 Distribution of aromatase in the brain of the African cichlid fish Astatotilapia burtoni: Aromatase expression, but not estrogen receptors, varies with female reproductive-state. J. Comp. Neurol. 528, 2499–2522. (doi:10.1002/cne.24908)

24. Nishiike Y et al. 2021 Estrogen receptor 2b is the major determinant of sex-typical mating behavior and sexual preference in medaka. Curr. Biol. 31, 1699–1710.e6. (doi:10.1016/j.cub.2021.01.089)

25. Koyama Y, Satou M, Oka Y, Ueda K. 1984 Involvement of the telencephalic hemispheres and the preoptic area in sexual behavior of the male goldfish, Carassius auratus: a brain-lesion study. Behav. Neural Biol. 40, 70–86. (doi:10.1016/S0163-1047(84)90182-1)

26. Kyle AL, Peter RE. 1982 Effects of forebrain lesions on spawning behaviour in the male goldfish. Physiol. Behav. 28, 1103–1109. (doi:10.1016/0031-9384(82)90183-4)

27. Satou M, Oka Y, Kusunoki M, Matsushima T, Kato M, Fujita I, Ueda K. 1984 Telencephalic and preoptic areas integrate sexual behavior in hime salmon (landlocked red salmon, Oncorhynchus nerka): Results of electrical brain stimulation experiments. Physiol. Behav. 33, 441–447. (doi:10.1016/0031-9384(84)90167-7)

28. Korzan WJ, Fernald RD. 2007 Territorial male color predicts agonistic behavior of conspecifics in a color polymorphic species. Behav. Ecol. 18, 318–323. (doi:10.1093/beheco/arl093)

29. Korzan WJ, Robison RR, Zhao S, Fernald RD. 2008 Color change as a potential behavioral strategy. Horm. Behav. 54, 463–470. (doi:10.1016/j.yhbeh.2008.05.006)

30. Clement TS, Grens KE, Fernald RD. 2005 Female affiliative preference depends on reproductive state in the African cichlid fish, Astatotilapia burtoni. Behav. Ecol. 16, 83–88. (doi:10.1093/beheco/arh134)

31. Renn SCP, Fraser EJ, Aubin-Horth N, Trainor BC, Hofmann HA. 2012 Females of an African cichlid fish display male-typical social dominance behavior and elevated androgens in the absence of males. Horm. Behav. 61, 496–503. (doi:10.1016/j.yhbeh.2012.01.006)

32. Jackson L, Dumitrascu M, Alward BA. 2023 Sex differences in aggression and its neural substrate in a cichlid fish. bioRxiv (doi:10.1101/2023.10.18.562975)

33. Maruska KP, Fernald RD. 2012 Contextual chemosensory urine signaling in an African cichlid fish., 68–74. (doi:10.1242/jeb.062794)

34. Field KE, Maruska KP. 2017 Context-dependent chemosensory signaling, aggression and neural activation patterns in gravid female African cichlid fish. J. Exp. Biol. 220, 4689–4702. (doi:10.1242/jeb.164574)

