## Supplementary Material for "Female cichlids attack and avoid—but will still mate with—androgen receptor mutant males that lack male-typical body coloration"

**Affiliations:** [1]University of Houston, Department of Psychology; [2]University of Houston, Department of Biology and Biochemistry; [3] These authors share first authorship.

**Supplementary Material**

**Supplementary Figures**

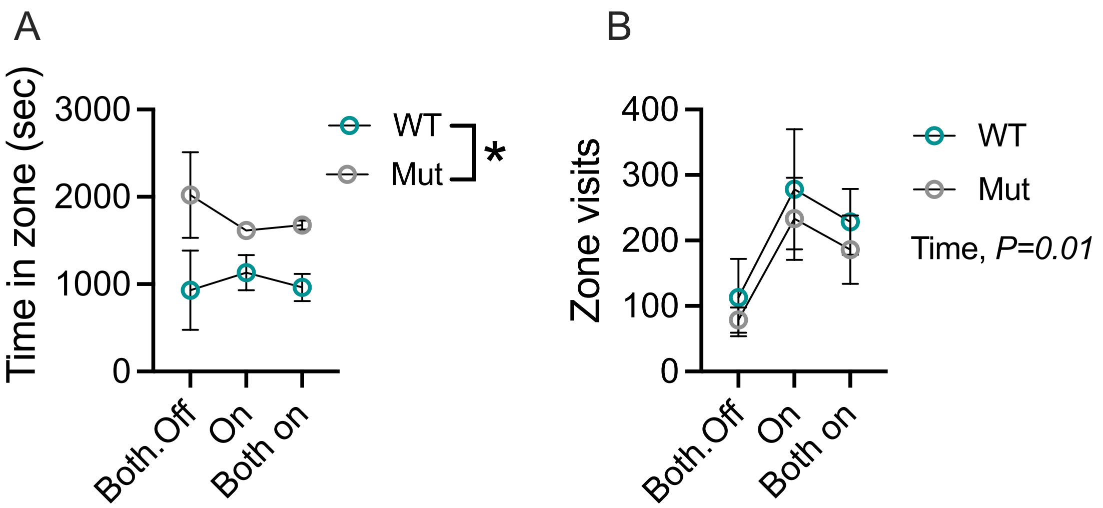


**Figure S1.** A) Time spent in female zone and B) zone visits by WT and Mut males during smartglass assays. “On” refers to the time during which only the WT or Mut smartglass was on and the other smartglass was off. Circles represent the Mean±Standard Error of the Mean (SEM). *=$P<0.05$.

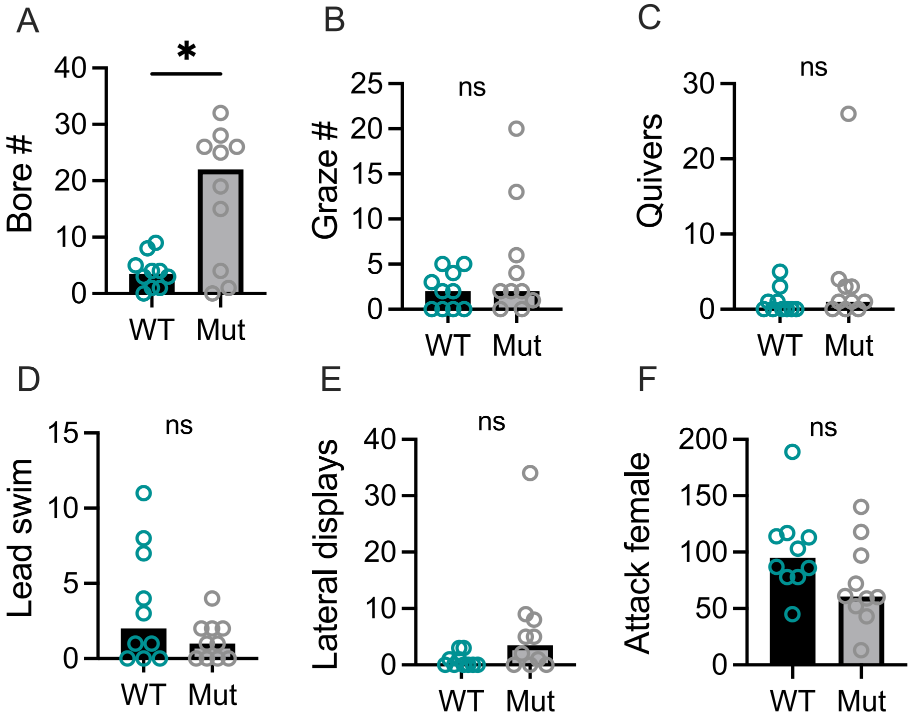


**Figure S2.** A-F) Behaviors performed by WT or Mut males when both smartglasses were on. Circles represent individual males during a given trial. Bars are the mean. *=*P<0.05.*

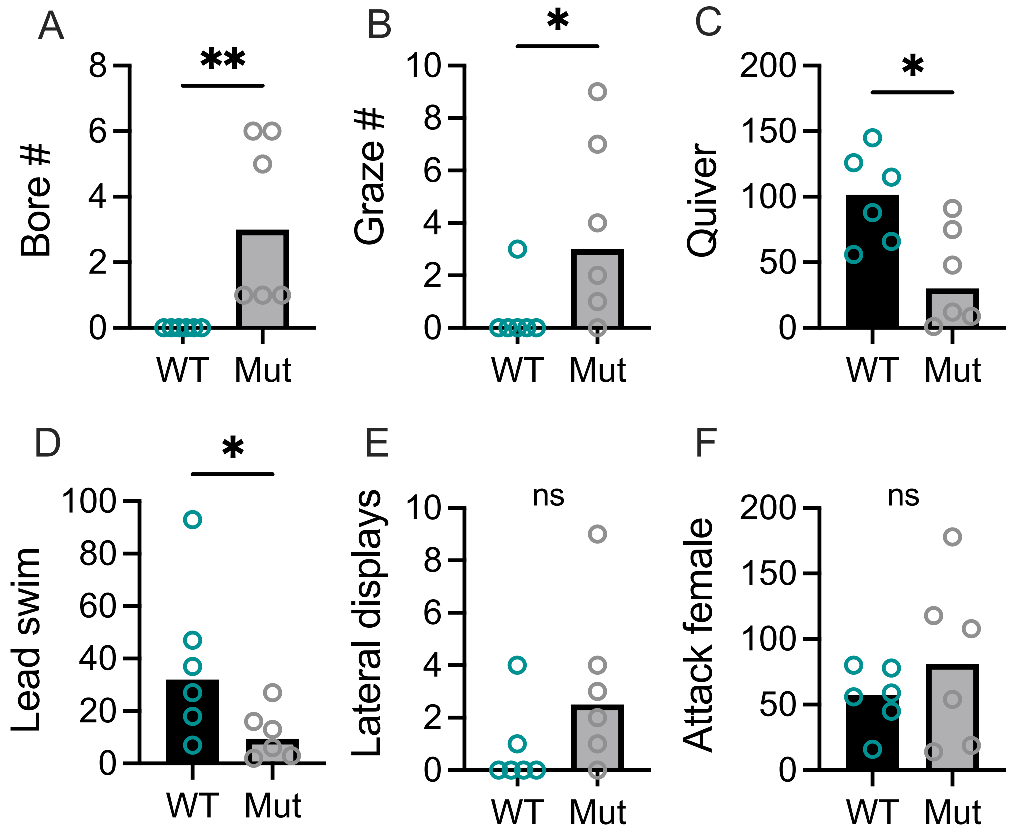


**Figure S3.** A-F) Behaviors performed by WT or Mut males during holistic assays. Circles represent individual males during a given trial. Bars are the mean. **=P<0.05. **=P<0.01.*

**Supplementary Videos**

**Supplementary Video Legends**

**Supplementary Video S1.** Example of female spawning with a WT male. This recording is of a holistic trial in which a female spawns with a WT male. Male and female are denoted during the video. A text indicator also appears when spawning starts in the video.

**Supplementary Video S2.** Example of female spawning with a Mut male. This recording is of a holistic trial in which a female spawns with a Mut male. Male and female are denoted during the video. A text indicator also appears when spawning starts in the video.
